# Ethylene-triggered subcellular trafficking of CTR1 enhances the response to ethylene gas

**DOI:** 10.1101/2020.06.12.148858

**Authors:** Han Yong Lee, Dong Hye Seo, Hye Lin Park, Arkadipta Bakshi, Chanung Park, Joseph J. Kieber, Brad M. Binder, Gyeong Mee Yoon

**Affiliations:** Department of Botany and Plant Pathology, Purdue University, West Lafayette, Indiana, 47907, USA; Center for Plant Biology, Purdue University, West Lafayette, Indiana, 47907, USA; Department of Biochemistry & Cellular and Molecular Biology, University of Tennessee,Knoxville, Tennessee, 37996, USA; Department of Biology, University of North Carolina, Chapel Hill, North Carolina, 27599, USA

**Keywords:** Ethylene, CTR1, Nuclear localization, Spatiotemporal regulation, Ethylene response growth kinetics

## Abstract

Ethylene gas controls plant growth and stress responses. Ethylene-exposed dark-grown seedlings exhibit dramatic growth reduction, yet the seedlings rapidly return to the basal growth rate when ethylene gas is removed. However, the underlying mechanism governing this reversible acclimation of dark-grown seedlings to ethylene remains enigmatic. Here, we report that ethylene triggers the translocation of the Raf-like protein kinase CONSTITUTIVE TRIPLE RESPONSE1 (CTR1), a negative regulator of ethylene signaling, from the endoplasmic reticulum to the nucleus. Nuclear-localized CTR1 stabilizes the ETHYLENE-INSENSITIVE3 (EIN3) transcription factor via interaction with the EIN3-BINDING F-box (EBF) proteins, thus enhancing the ethylene response and delaying growth recovery. These findings uncover a mechanism of the ethylene signaling pathway that links the spatiotemporal dynamics of cellular signaling components to organismal responses.

## Introduction

The ability of organisms to respond to and integrate environmental signals leading to an appropriate response is critical for optimal growth and development, particularly for plants, which are non-motile. Plants adapt to a wide variety of abiotic stresses, and after removal of stress, they need to rapidly restore basal cellular homeostasis. One key signal for the acclimation of plants to abiotic stress is the plant hormone ethylene. Ethylene regulates multiple aspects of growth and development, including fruit ripening, leaf and floral senescence, cell elongation, seed germination, root hair formation, and responses to biotic and abiotic stress^1–3^. Ethylene-mediated stress acclimation includes, but is not limited to, the rapid elongation of rice internodes in response to flooding, drought responses, salt tolerance, heavy metal tolerance, and morphological changes in roots in response to nutrient deficiency^4–8^.

Extensive molecular genetic studies have elucidated the basic ethylene signaling pathway^2, 9–12^. In the absence of ethylene, the endoplasmic reticulum (ER)-localized ethylene receptors activate CONSTITUTIVE TRIPLE RESPONSE1 (CTR1) protein kinase, which in turn phosphorylates Ethylene-Insensitive 2 (EIN2), an ER membrane-localized Nramp homolog that positively regulates ethylene responses. CTR1-mediated phosphorylation of EIN2 results in preventing EIN2 from signaling in the absence of ethylene^13–15^. In response to ethylene, the receptors and hence CTR1 are inactivated, leading to reduced phosphorylation and increased accumulation of EIN2. EIN2 is then proteolytically cleaved, and the C-terminal domain (EIN2-CEND) is released. EIN2-CEND then translocates into the nucleus and indirectly activates the Ethylene-Insensitive 3 (EIN3) and EIN3-like (EIL) paralogs, which are central transcription factors in ethylene signaling^13–15^. EIN2-CEND also associates with the mRNAs of EIN3-Binding F-box 1 (*EBF1*) and *EBF2*, and subsequently represses their translation, thus ultimately blocking the degradation of EIN3/EIL proteins ^16, 17^. The function of CTR1 beyond phosphorylating EIN2 at the ER, if any, has not been characterized.

The hypocotyls of dark-grown seedlings exposed to ethylene show a dramatically diminished growth rate. However, after the removal of ethylene, the seedlings return to a basal growth rate within 90 min^18, 19^, although the proteolytic cleavage of EIN2 is irreversible. How this fast growth recovery is regulated remains an open question. Here, we report that CTR1 rapidly translocates from the ER to the nucleus in response to ethylene. Unexpectedly, the nuclear-localized CTR1 enhances ethylene responses through a mechanism that does not require its kinase activity and inhibits the fast recovery of seedling growth back to basal levels. These results suggest a new mechanistic paradigm for the dynamic regulation of the ethylene signaling involving the translocation of CTR1 to the nucleus, thereby strengthening EIN2-mediated EIN3 activation in the nucleus.

## Results

### Ethylene-induced ER-to-nucleus translocation of CTR1

CTR1 consists of an N-terminal regulatory domain and a C-terminal kinase domain that is homologous to the catalytic domain of the Raf kinase family and lacks any organelle targeting sequences, including canonical nuclear localization sequences (NLSs)^11^ (Fig. 1a). To determine the role of CTR1 beyond its regulation of EIN2, we examined the subcellular localization of CTR1 after plant exposure to exogenous ethylene in the dark. To this end, we introduced a ~7.6 Kb genomic *CTR1* transgene containing a *CTR1* promoter region (0.7 Kb upstream of the 5’UTR) driving expression of the CTR1 coding region fused to a GFP reporter (*CTR1p:GFP-gCTR1*)^11^. The *CTR1p:GFP-gCTR1* transgene fully complemented the *ctr1-2* phenotypes in both light- and dark-grown seedlings, including decreased rosette and inflorescence size, and a strong triple response phenotype (Fig. 1b and Supplementary Fig. 1), indicating that the fusion protein was functional. In the presence of an inhibitor of ethylene perception, silver nitrate (AgNO_3_), or in an *etr1-1* mutant background, the GFP-CTR1 fusion protein localized to the ER (Fig. 1c, d), in agreement with findings from previous reports^20, 21^. However, interestingly, in response to either 1-aminocyclopropane-1-carboxylate (ACC, a direct precursor of ethylene) or exogenous ethylene, GFP-CTR1 accumulated in the nucleus (Fig. 1c, 1e, and 1f). Disruption of either EIN2 or EIN3/EIL1 did not prevent the nuclear accumulation of CTR1 (Fig. 1g), suggesting that EIN2 and EIN3 are not required for CTR1 nuclear translocation. Furthermore, when GFP-CTR1 was expressed under the control of the strong CaMV 35S promoter, a large fraction of GFP-CTR1 constitutively localized to the nucleus in etiolated seedlings, regardless of ACC treatment (Fig. 1h and Supplementary Fig. 2), presumably because of the limited number of ethylene receptors tethering CTR1 to the ER. Fractionation analysis confirmed that CTR1 was enriched in the nuclear fraction in extracts of ACC-treated seedlings, but remained soluble in extracts derived from seedlings grown in the presence of AgNO_3_ (Fig. 1i). Fluorescence recovery after photobleaching analysis showed that the movement of CTR1 into the nucleus was visible at 15 sec after photobleaching of the entire nucleus, and GFP-CTR1 fluorescence in the nucleus recovered by 40% at 147 sec after photobleaching (Supplementary Fig. 3). This result further demonstrated that CTR1 migrates into the nucleus, and the nuclear-localization of CTR1 does not result from increased stability of preexisting CTR1 in the nucleus. Because both ACC and ethylene caused equivalent CTR1 nuclear translocation (Fig. 1e), we used ACC for further studies.

**Fig. 1.**
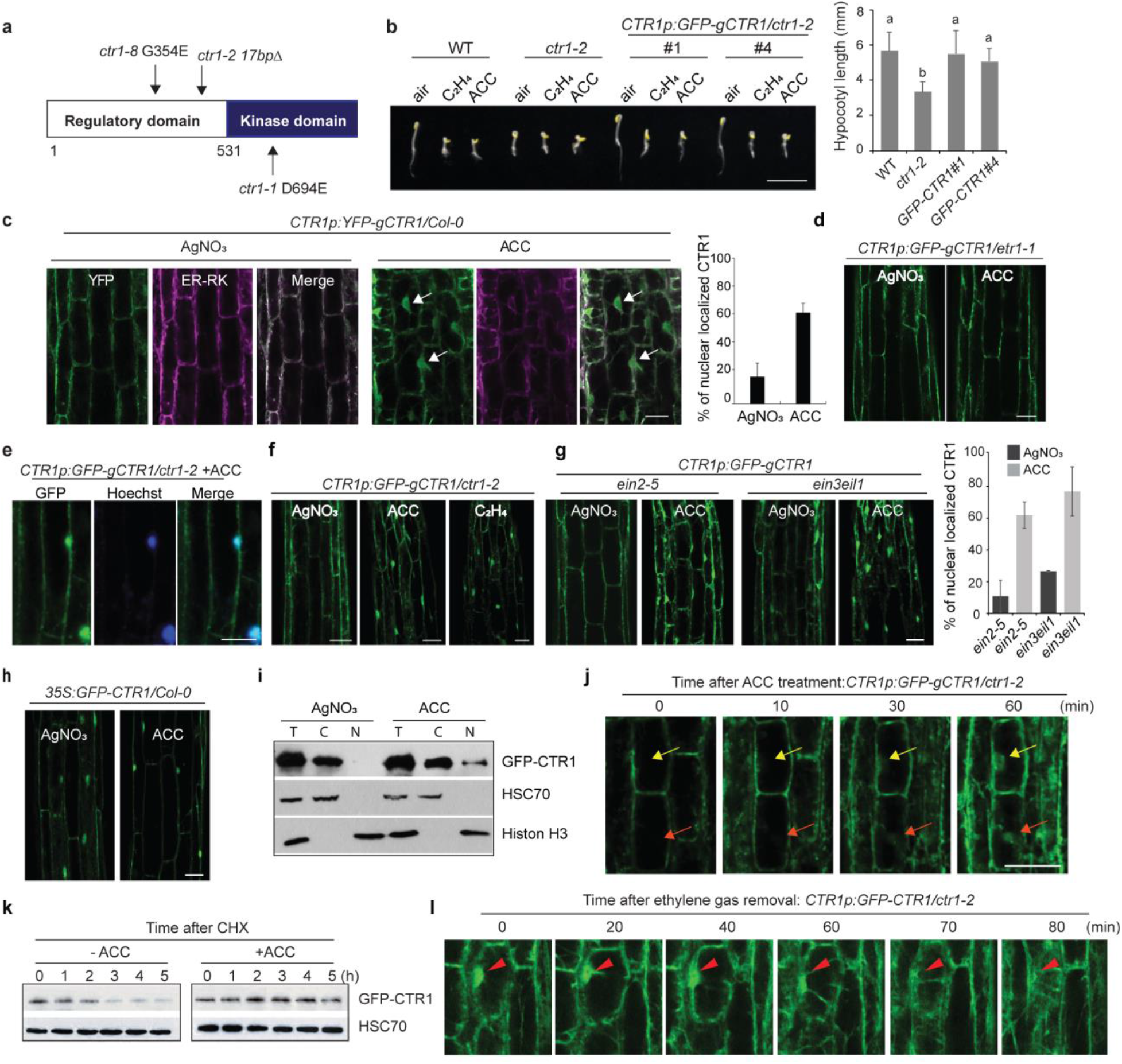
Ethylene-activated CTR1 translocation to the nucleus does not require EIN2 and EIN3. **a,** Diagram of the CTR1 protein domain structure. The positions of *ctr1* mutant alleles are indicated by arrows. **b**, The GFP-fused WT genomic *CTR1* fragment fully rescues *ctr1-2* and confers an ethylene response. Seedlings were grown for 3 d in the dark with or without ACC or ethylene. The graph represents the quantification of hypocotyl lengths of seedlings grown on MS without ACC. MS, Murashige and Skoog medium. Different letters indicate significant differences at *p* < 0.05 (one-way ANOVA, post-hoc Tukey’s HSD). Data represent the means and SD (*n* ≥ 19). Scale bar, 5 mm. **c,** Three-day-old etiolated seedlings co-expressing GFP-CTR1 and ER-RK were grown on MS medium with AgNO_3_ or treated with ACC for 2 h. The areas below the hook and above the elongation zone were imaged. ER-RK^41^, an mCherry-fused ER marker. Arrows indicate nuclear-localized GFP-CTR1. The graph represents the ratio of nuclear-localized CTR1 in dark-grown seedlings. Error bars indicate SD (*n* ≥ 3). **d**, GFP-CTR1 does not translocate into the nucleus in *etr1-1* mutant background. **e**, Ethylene and ACC activate CTR1 nuclear translocation. Seedlings were grown on MS medium with or without 100 μM AgNO_3_ in the dark. Seedlings grown on MS without AgNO_3_ were treated with 200 μM ACC for 2 h. For ethylene treatment, seedlings were grown in vials without capping for 3 d, followed by capping and injection of 10 ppm ethylene for 2 h before imaging. Scale bars, 50 μm. **f**, GFP-CTR1 fluorescence overlaps with Hoechst nuclear staining under ACC treatment, showing CTR1 nuclear localization. **g,** Seedlings expressing GFP-CTR1 in *ein2-5, ein3 eil1*, and *etr2-3ers2-3ein4-4* mutants were grown on MS medium with or without AgNO_3_ for 3 d. For ACC treatment, the seedlings grown on MS medium without AgNO_3_ were treated with ACC for 2 h before visualization by confocal microscopy. The graph represents the ratio of nuclear-localized CTR1 in dark-grown seedlings. Error bars indicate SD (*n* ≥ 3). **h,** Overexpression of CTR1 from 35S promoter leads to constitutive nuclear localization of CTR1 in dark-grown seedlings. **i**, The nuclear (N) and cytoplasmic (C) distribution of CTR1 in seedlings with or without ACC treatment. The etiolated *CTR1p:GFP-gCTR1/ctr1-2* seedlings were treated with ACC (200 μM) or were grown on medium containing AgNO_3_ (100 μM) for 3 d in the dark. Total protein extract (T) was fractionated into nuclear and cytoplasmic fractions, then subjected to immunoblotting analysis with anti-GFP, anti-Histone H3, and anti-HSC70. **j,** Time-lapse image series of hypocotyl cells expressing GFP-CTR1 in 3-d-old etiolated seedlings after exposure to 200 μM ACC, visualized by confocal microscopy. Arrows track specific cell nuclei, showing the accumulation of GFP-CTR1 in response to ACC. **k**, Protein degradation analysis of CTR1. *CTR1p:GFP-gCTR1/ctr1-2* seedlings were treated with or without 10 μM ACC. The seedlings were subsequently treated with 250 μM cycloheximide (CHX), and total protein extracts were used for immunoblotting. **l,** Time-lapse image analysis of hypocotyl cells expressing GFP-CTR1. The disappearance kinetics of nuclear-localized GFP-CTR1 was observed in the seedlings pretreated with 10 ppm ethylene. Arrows track specific cell nuclei, showing decreased nuclear-localized CTR1 after ethylene purging. All scale bars represent 50 μm, except the scale bar in (**b**). Ten or more seedlings from at least two independent lines were observed for CTR1 localization, and representative images are shown. The areas below the hook and above the elongation zone of the hypocotyls of dark-grown seedlings were imaged.

EIN2-CEND migrates into the nucleus within 10 min after ethylene treatment^14^. We observed comparable kinetics of EIN2 nuclear movement in response to ACC (Supplementary Fig. 4). We monitored the dynamics of CTR1 movement in response to increasing ACC treatment duration (Fig. 1j). CTR1 first accumulated to detectable levels in the nucleus 30 min after ACC treatment, and a further increase in nuclear protein levels was observed after an additional 30 min (Fig. 1j). The overall GFP fluorescence intensity was enhanced with longer ACC exposure, consistent with increased stabilization of the CTR1 protein in response to ACC (Fig. 1k), similarly to previously reported findings^22^. The GFP fluorescence of nuclear GFP-CTR1 began to diminish within 60 min after ethylene removal, and by 80 min after ethylene removal, no detectable fluorescence remained in the nucleus (Fig. 1l). Together, these findings revealed that ethylene stimulates the translocation of CTR1 from the ER to the nucleus in an EIN2 and EIN3/EIL independent manner, and that nuclear CTR1 protein may undergo degradation via an unknown mechanism or may be exported back to the cytoplasm after ethylene removal.

### The CTR1 N-terminus inhibits CTR1 nuclear trafficking

CTR1 is a peripheral membrane protein, which is recruited to the ER via interaction with ethylene receptors, where it acts to prevent ethylene signaling by phosphorylation of EIN2^11, 13, 22^. Nuclear translocation of CTR1 requires the dissociation of CTR1 from the ER and presumably disassociation from the ethylene receptors. CTR1 interacts with the ethylene receptor via its N-terminal domain^20^. Therefore, we examined whether the N-terminal domain of CTR1 might affect its nuclear translocation. To test this possibility, we expressed a fusion protein of the CTR1 lacking the N-terminal domain (ΔNT-CTR1) (Fig. 1a) from its native promoter in stable transgenic plants. The *CTR1p:GFP-ΔNT-gCTR1* transgene did not rescue *ctr1-2* in either etiolated or light-grown plants (Fig. 2a and Supplementary Fig. 1). Consistent with this finding, *CTR1p:GFP*-Δ*NT-gCTR1* seedlings showed constitutive expression of the ethylene-inducible *ERF1* gene, to a level comparable to that observed in *ctr1-2* (Fig. 2b). The failure of the Δ*NT-gCTR1* transgene to complement *ctr1-2* probably resulted from the decreased ER membrane targeting of the ΔNT-CTR1 protein. Indeed, the ΔNT-CTR1 protein was constitutively localized to the nucleus in the dark-grown seedlings, whether expressed from its own promoter or the CaMV 35S promoter (Fig. 2c and 2d).

**Fig. 2.**
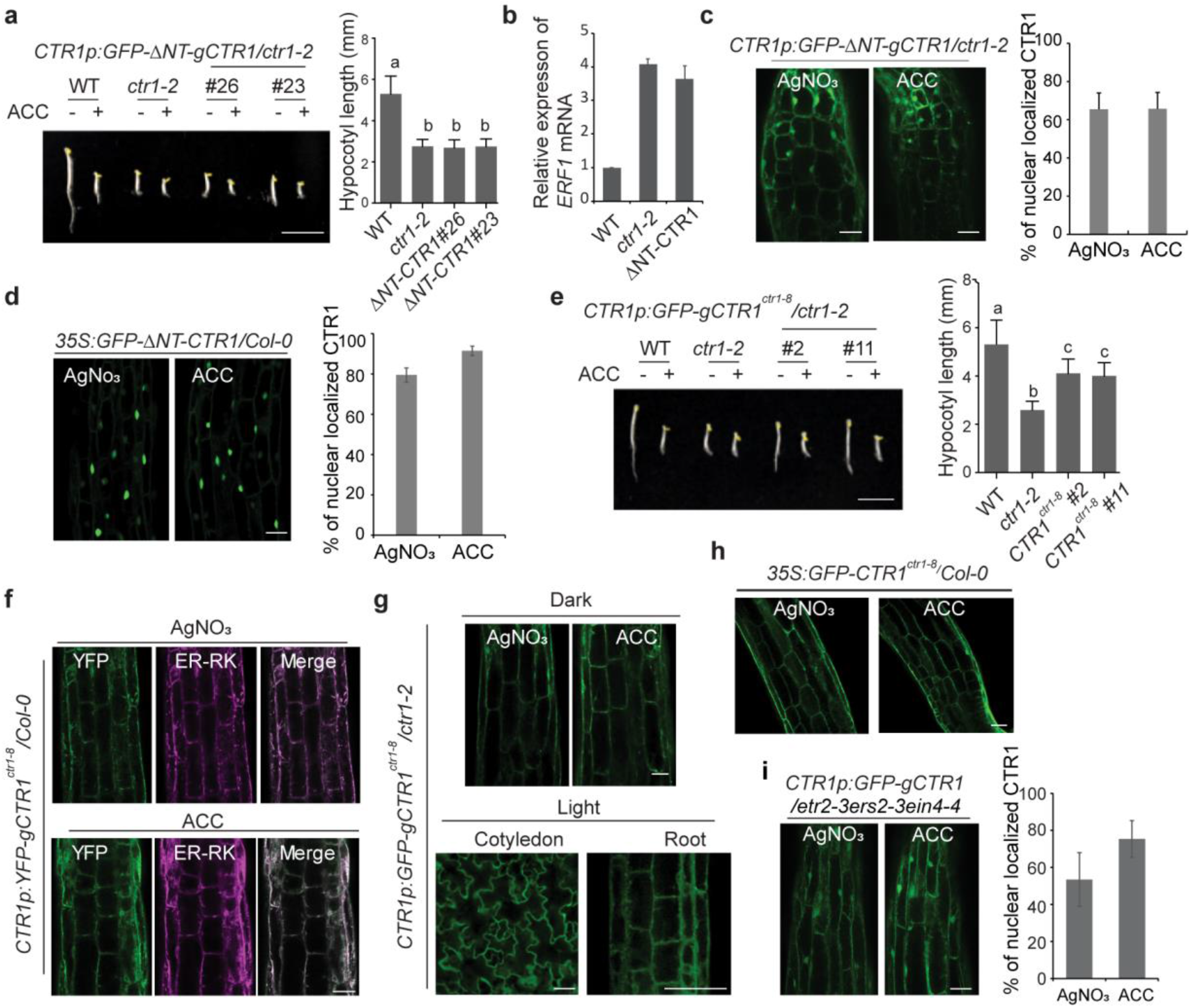
The N-terminus of CTR1 inhibits ACC-induced CTR1 nuclear movement. **a,** Seedlings were grown on MS medium with or without 10 μM ACC for 3 d and photographed. The graph shows quantification of the hypocotyl lengths of seedlings grown on MS without ACC. Scale bar, 5 mm. Different letters indicate significant differences at *p* < 0.05 (one-way ANOVA, post-hoc Tukey’s HSD). Data represent the means and SD (*n* ≥ 43). **b,** Quantitative gene expression analysis of ethylene-responsive *ERF1* in WT, *ctr1-2* and *CTR1p:GFP-ΔNT-gCTR1* seedlings without ACC treatment. Expression was normalized to an *Actin* control and is presented relative to the untreated WT control. Data represent the means and SD for three biological replicates. **c**, Constitutive nuclear localization of GFP-ΔNT-CTR1 expressed from the native promoter in the hypocotyls of dark-grown seedlings. The graph represents the ratio of nuclear-localized CTR1 in dark-grown seedlings treated with ACC or AgNO_3_. Error bars indicate SD (*n* ≥ 3). **d**, Constitutive nuclear localization of GFP-ΔNT-CTR1 expressed from the CaMV 35S promoter in 3-d-old dark-grown. The graph represents the ratio of nuclear-localized CTR1 in dark-grown seedlings treated with ACC or AgNO_3_. Error bars indicate SD (*n* ≥ 3). **e**, Seedlings expressing GFP-CTR1^ctr1-8^ from the native promoter were grown on MS medium with or without 10 μM ACC for 3 d and photographed. The graph represents the quantification of hypocotyl lengths of seedlings grown on MS without ACC. Scale bar, 5 mm. Different letters indicate significant differences at *p* < 0.001 (one-way ANOVA, post-hoc Tukey’s HSD). Data represent the means and SD (*n* ≥ 37). **f**, Seedlings co-expressing YFP-CTR1^ctr1-8^ and ER-RK were grown in the dark and used to visualize co-localization. **g-h**, GFP-CTR1^ctr1-8^ does not translocate into the nucleus in both dark and light conditions. **I,** Constitutive nuclear localization of GFP-CTR1 in the *etr2-3ers2-3ein4-4* mutant background in the dark-grown seedlings. Scale bars in c, d, f, g, h, and i represent 50 μm. Ten or more seedlings from at least two independent lines were observed for CTR1 localization, and representative images are shown. The areas below the hook and above the elongation zone of the hypocotyls of dark-grown seedlings were used for imaging.

We further explored whether binding of CTR1 to the ethylene receptors might play a role in its nuclear localization. Previous studies have shown that the *ctr1-8* mutation (G354E) blocks the interaction of CTR1 with ETR1 in yeast-two-hybrid assays but does not affect the intrinsic kinase activity of the protein (Fig. 1a)^23^. In agreement with the weak hypermorphic nature of *ctr1-8*, the *GFP-CTR1^ctr1-8^* transgene partially complemented *ctr1-2* in both light- and dark-grown seedlings (Fig. 2e and Supplementary Fig. 1). Previous fractionation studies have demonstrated that the *ctr1-8* mutant protein is found primarily in the soluble fraction, and a minor portion remains associated with the membrane fraction^21^, in contrast to the predominant ER localization of wild-type (WT) CTR1. However, we observed that a large fraction of GFP-CTR1^ctr1-8^ still appeared to localize to the ER (Fig. 2f). We speculate that this finding might have been due to weak association of the CTR1^ctr1-8^ protein with the receptors, thus resulting in a rapid equilibrium being reached between the receptor-bound and unbound states of CTR1^ctr1-8^. Unexpectedly, GFP-CTR1^ctr1-8^ did not translocate to the nucleus in either etiolated or light-grown seedlings, regardless of ACC treatment (Fig. 2f, 2g and 2h). To further examine the role of CTRI’s interaction with ethylene receptors in regulating its nuclear translocation, we examined the localization of GFP-CTR1 in various ethylene receptor mutant backgrounds. GFP-CTR1 constitutively localized to the nucleus in the *etr2-3ers2-3ein4-4* triple mutants (Fig. 2i), in which three of the five ethylene receptors were disrupted, as well as in additional multiple loss-of-function receptor mutants (Supplemental Fig. 5). These results suggested that in the absence of ethylene, ethylene receptors are likely to tether CTR1 at the ER via direct interaction, thus preventing CTR1 nuclear translocation.

### The nuclear movement of CTR1 is independent of kinase activity

To address the role of kinase activity in CTR1 localization, we expressed a catalytically inactive *ctrl* mutant (*CTR1p:GFP-gCTR1^ctr1-1^*) in the *ctr1-2* mutant background (Fig. 1a)^23^. The *GFP-CTR1^ctr1-1^* transgene did not rescue *ctr1-2* in either the dark or light conditions (Fig. 3a and Supplementary Fig. 1) consistent with its strongly hypomorphic nature. Similar to the WT CTR1, full-length inactive GFP-CTR1^*ctr1-1*^ expressed from its native promoter translocated to the nucleus after ACC treatment (Fig. 3b). When expressed from the strong CaMV 35S promoter, GFP-CTR1^*ctr1-1*^ constitutively localized in the nucleus, similarly to WT CTR1 (Fig. 3c). In addition, inactive ΔNT-CTR1^*ctr1-1*^ expressed from either the CaMV 35S or the native promoter (*CTR1p:GFP*-Δ*NT-gCTR1/ctr1-2* and *35S:GFP*-Δ*NT-CTR1^ctr1^/Col-0*) constitutively localized to the nucleus (Fig. 3d and 3e), and this Δ*NT-CTR^ctr1-1^* transgene did not rescue *ctr1-2* in either dark- or light-grown seedlings (Supplementary Fig. 1 and 6).

**Fig. 3.**
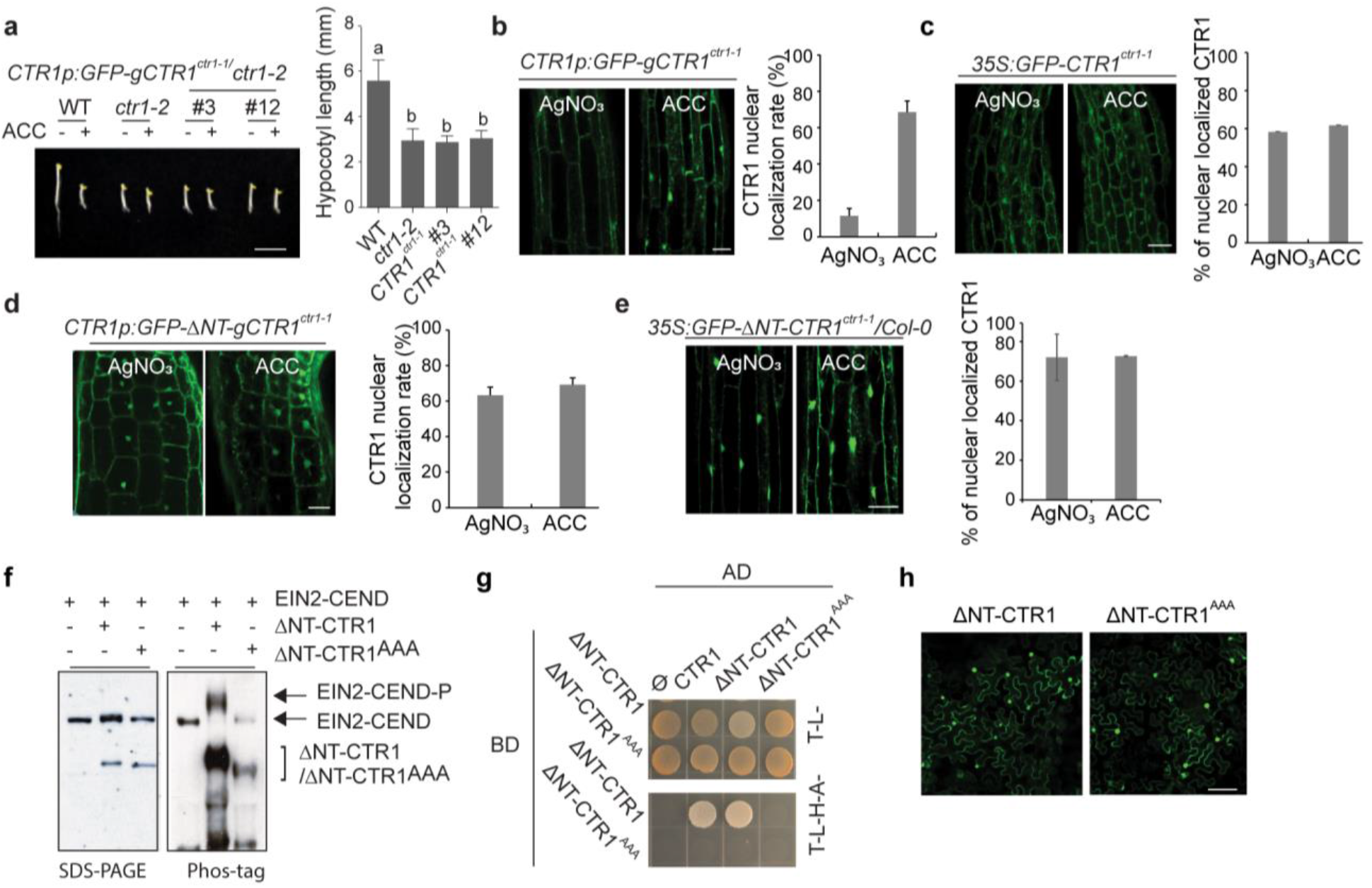
The kinase activity of CTR1 is not necessary for ACC-induced CTR1 nuclear translocation. **a**, Seedlings were grown on MS medium with or without 10 μM ACC for 3 d and photographed. Scale bar, 5 mm. The graph shows the quantification of hypocotyl lengths of seedlings grown on MS without ACC. Different letters indicate significant differences at *p* <0.05 (one-way ANOVA, post-hoc Tukey’s HSD). Data represent the means and SD (*n* ≥ 43). **b-c**, Seedlings expressing GFP-CTR1^ctr1-1^ from its native promoter (**b**) or the CaMV 35S promoter (**c**) were grown on MS medium with or without AgNO_3_. For ACC treatment, the seedlings grown on MS medium without AgNO_3_ were further treated with ACC for 2 h before imaging. The graph represents the ratio of nuclear-localized CTR1 in dark-grown seedlings treated with ACC or AgNO_3_. Error bars indicate SD (*n* ≥ 3). **d-e**, Seedlings expressing GFP-ΔNT-CTR1^ctr1-1^ from its native promoter (**d**) or the CaMV 35S promoter (**e**) were grown on MS medium with or without AgNO_3_. The graph represents the ratio of nuclear-localized CTR1 in dark-grown seedlings treated with ACC or AgNO_3_. Error bars indicate SD (*n* ≥ 3). **f**, *In vitro* phos-tag analysis of active and inactive ΔNT-CTR1 in *Arabidopsis* protoplasts. **g**, The indicated bait and prey constructs were co-transformed into the AH109 yeast strain, and the transformed yeast were grown on selection medium. **h**, Tobacco leaves were infiltrated with agrobacterium transformed with ΔNT-CTR1 or ΔNT-CTR1^AAA^ plasmid construct, followed by 3 d incubation and visualization of nuclear signals by confocal microscopy. All scale bars represent 50 μm except the scale bar in (**a**). Ten or more seedlings from at least two independent lines were observed for CTR1 localization, and representative images are shown. The areas below the hook and above the elongation zone of hypocotyls of dark-grown seedlings were used for imaging.

ΔNT-CTR1 is autophosphorylated on four residues (S703/T704/S707/S710) within the kinase activation loop, and this autophosphorylation is critical for CTR1 kinase activity and homodimer formation^24^. To test the role of this autophosphorylation, we altered three of these target S/T residues (T704/S707/S710) to Ala (CTR1^AAA^), which has been shown to disrupt homodimer formation^24^. We confirmed that ΔNT-CTR1^AAA^ was catalytically inactive toward the EIN2 substrate; WT ΔNT-CTR1 but not ΔNT-CTR1AAA phosphorylated EIN2-CEND when co-expressed in *Arabidopsis* mesophyll protoplasts (Fig. 3f). Whereas WT ΔNT-CTR1 interacted with itself in a yeast-two-hybrid assay, the ΔNT-CTR1AAA did not interact with itself (Fig. 3g). Similar to WT ΔNT-CTR1, ΔNT-CTR1-KD^AAA^ constitutively localized to the nucleus (Fig. 3h). Together, these results indicated that kinase activity and probably homodimerization are not required for CTR1 nuclear translocation.

### Nuclear-localized CTR1 delays growth recovery of seedlings after ethylene-induced growth inhibition

Both the *ctr1-1* and *ctr1-8* mutants show constitutive ethylene responses^23^, despite that the CTR1^*ctr1-1*^ and CTR1^*ctr1-8*^ proteins showed different nuclear translocation responses to ethylene (Fig. 2f, 2g, 2h, and 3b). This finding indicated that CTR1 nuclear movement does not control the primary ethylene response but instead may influence ethylene response kinetics by fine-tuning nuclear ethylene signaling. To test this hypothesis, we measured the ethylene growth response kinetics of the hypocotyls of etiolated seedlings, which have been widely exploited to analyze ethylene mutants^18, 25^. There are two phases of growth inhibition of WT *Arabidopsis* hypocotyls in response to ethylene. Phase I begins 10 min after ethylene treatment and is characterized by a rapid deceleration in growth rate. After a transient (15 min) plateau in the growth rate, phase II growth inhibition is initiated, thus resulting in further growth suppression lasting 30 min until the growth rate reaches a new low steady-state rate^18, 25^. Genetic studies have revealed that EIN2 is necessary for both phases, but only phase II requires EIN3/EIL1^26^. Interestingly, after removal of ethylene during phase II, hypocotyl growth rapidly recovers to the pre-treatment growth rate within 90 min^18, 25^, indicating the existence of a mechanism to rapidly shut off the ethylene response.

To examine whether nuclear-localized CTR1 might play a role in the rapid growth inhibition when ethylene is added or in the recovery kinetics when ethylene is removed, we performed time-lapse analyses of the ethylene response growth kinetics of seedlings overexpressing GFP-ΔNT-CTR1 (*35S:GFP*-Δ*NT-CTR1/Col-0*) and GFP-ΔNT-CTR1^ctr1-1^ (35S: *GFP*-Δ*NT-CTR1^ctr1-1^/Col-0*), both of which showed strong constitutive nuclear localization and exhibited an ethylene response in the dark (Fig. 2d and 3e). After ethylene exposure, both *35S:GFP*-Δ*NT-CTR1/Col-0* and *35S:GFP*-Δ*NT-CTR1^ctr1-1^/Col-0* seedlings had similar onset and strength of phase I and II growth inhibition to those in the WT seedlings. However, the recovery of the hypocotyl growth rate of both *35S:GFP*-Δ*NT-CTR1/Col-0* and *35S:GFP*-Δ*NT-CTR1^ctr1-1^/Col-0* seedlings after ethylene removal was slower (~30 min) than that of the WT (Fig. 4a). Similar kinetics of ethylene response and growth recovery was also observed in seedlings expressing active or inactive WT full-length CTR1 (*35S:GFP-CTR1/Col-0* and *35S:GFP-CTR1^ctr1-1^/Col-0*), both of which showed some level of constitutive CTR1 nuclear localization (Fig. 1h, 3c, and Supplementary Figure 7). The delayed hypocotyl growth recovery in *35S:GFP*-Δ*NT-CTR1/Col-0, 35S:GFP*-Δ*NT-CTR1^ctr1-1^/Col-0, 35S:GFP-CTR1/Col-0* and *35S:GFP-CTR1^ctr1-1^/Col-0* resembled that reported in previous studies with EIN3 overexpression or loss of EBF2^19^.

**Fig. 4.**
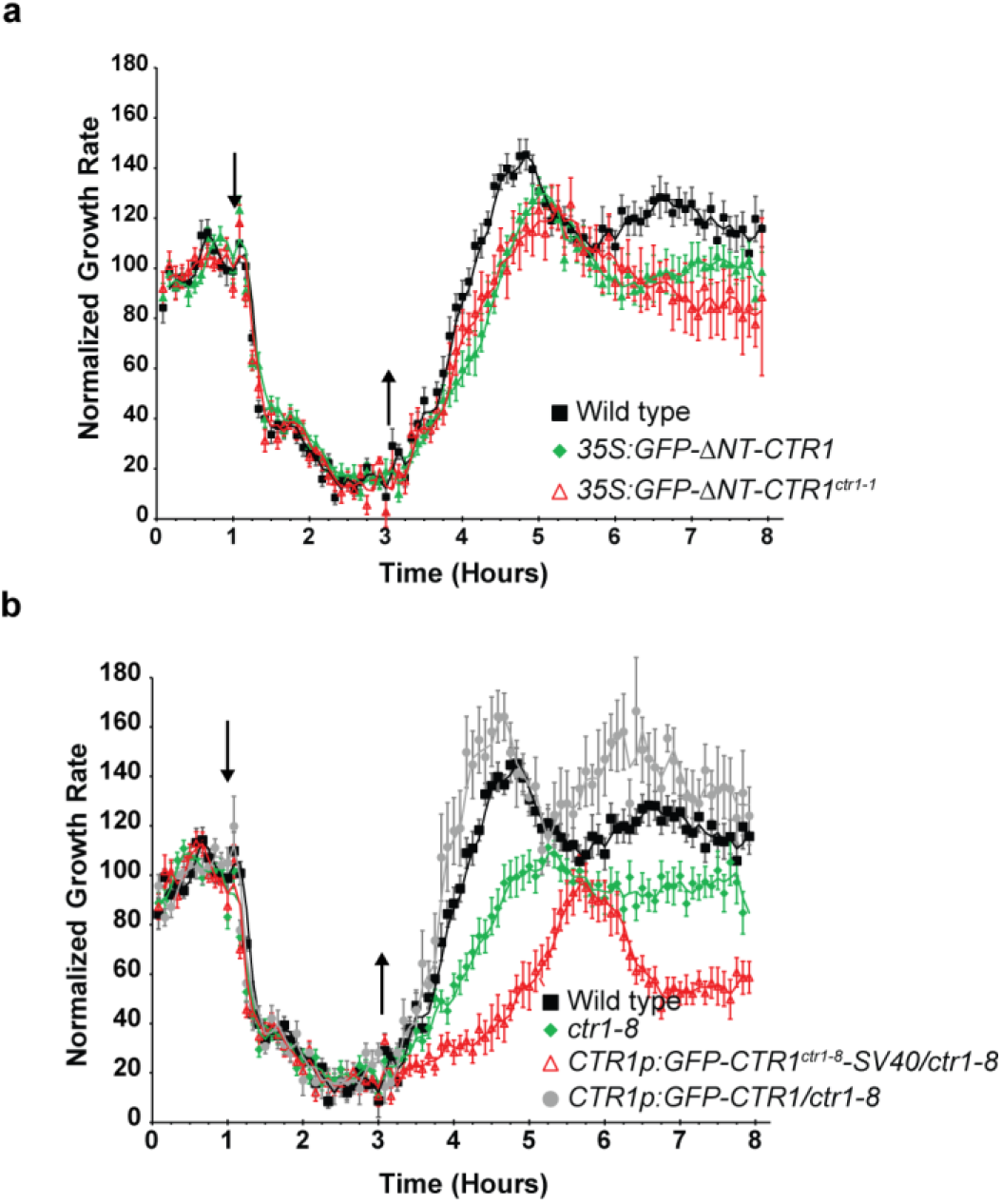
Nuclear-localized CTR1 delays growth recovery of hypocotyls after removal of ethylene. The hypocotyl growth rate in response to ethylene was recorded for 1 h in air, followed by 2 h exposure to 10 ppm ethylene and then 5 h recovery in air. Ethylene was introduced 1 h after measurements were initiated (down arrow) and then removed 2 h later (up arrow). The responses of WT seedlings are shown in each graph. **a,** *35S:GFP-ΔNT-CTR1* and *35:GFP-ΔNT-CTR1^ctr1-1^*. **b,** *ctr1-8, CTR1p:GFP-CTR1^ctr1-8^-SV40/ctr1-8*, and *CTR1p:GFP-CTR1/ctr1-8*. Data were normalized to the growth rate in air before treatment with ethylene. Error bars indicate SE (*n* ≥ 6). The experiments were repeated at least three times and generated similar results.

The *ctr1-8* mutant recovered approximately 90 min later than the WT after the removal of ethylene (Fig. 4b). This result was opposite from our expectations, because CTR1^ctr1-8^ protein does not translocate to the nucleus, unlike ΔNT-CTR1 and ΔNT-CTR1^ctr1-1^ proteins. However, given the hypermorphic nature of the mutation, the slower growth recovery of the *ctr1-8* mutant might be attributable to its weak interaction with the receptors, thus resulting in decreased EIN2 phosphorylation and consequently enhanced EIN3 levels. To further investigate the role of CTR1 in the recovery kinetics, we introduced a WT full-length genomic *CTR1* fragment into the *ctr1-8* mutant (*CTR1p:GFP-gCTR1/ctr1-8*). The *GFP-gCTR1* transgene rescued the phenotypes of *ctr1-8* in the dark (Supplementary Figure 8) and restored the slower growth recovery kinetics of *ctr1-8* to levels comparable to those of the WT, thus confirming that CTR1 is responsible for the slower recovery kinetics (Fig. 4b). Next, we examined the correlation between CTR1 nuclear translocation and ethylene growth recovery. To this end, we generated a transgenic line expressing simian virus (SV40) NLS-fused CTR1^ctr1-8^ protein in a *ctr1-8* mutant background (*CTR1p:GFP-gCTR1^ctr1-8^-SV40/ctr1-8*) to determine whether enhanced levels of nuclear-localized CTR1 might further delay the slower recovery kinetics of the *ctr1-8* mutant. The *GFP-gCTR1^ctr1-8^-SV40* transgene did not complement *ctr1-8* and exhibited a comparable ACC response to that of *ctr1-8* (Supplementary Figure 8). As expected, GFP-CTR1^ctr1-8^-SV40 proteins constitutively localized in the nucleus (Supplementary Figure 8). *CTR1p:GFP-gCTR1^ctr1-8^-SV40/ctr1-8* hypocotyls recovered approximately 45 min more slowly than *ctr1-8* after ethylene removal (Fig. 4b), thus indicating that nuclear-localized CTR1 promotes slower growth recovery. Together, these results suggest that nuclear-localized CTR1 delays the growth recovery of seedlings after ethylene removal, presumably via the stabilization of EIN3 proteins, without needing CTR1 kinase activity.

### Nuclear-localized CTR1 stabilizes EIN3 via EBFs without kinase activity

The correlation between nuclear-localized CTR1 and the delayed growth recovery kinetics suggests that CTR1 upregulates nuclear ethylene responses. Therefore, we explored whether CTR1 might modulate EIN3 function in the nucleus in an ethylene-dependent manner. To test this possibility, we first determined whether CTR1 interacts with ethylene signaling components in the nucleus, including EIN2-CEND, EIN3, and EBFs, by using yeast-two-hybrid and BiFC assays. We used CTR1 with an N-terminal deletion in the yeast-two hybrid assay because the full-length protein is autoactivated (Supplementary Fig. 9). Both ΔNT-CTR1 and ΔNT-CTR1^ctr1-1^ interacted with EIN2-CEND, EBF1, and EBF2, but not with EIN3 (Fig. 5a). Full-length CTR1 interacted with the EBFs in a BiFC assay regardless of ACC treatment (Fig. 5b and Supplementary Fig. 10), but not with EIN3 and EIN2-CEND. EIN2-CEND reconstituted YFP signals with its known nuclear interacting protein, EIN2 NUCLEAR-ASSOCIATED PROTEIN 1 (ENAP I) in the nucleus (Supplementary Fig. 11) ^27^, thus indicating that EIN2-CEND was expressed and present in the nucleus in these assays. Co-immunoprecipitation assays further confirmed that CTR1 interacts with EBF2 *in vivo* (Fig. 5c).

**Fig. 5.**
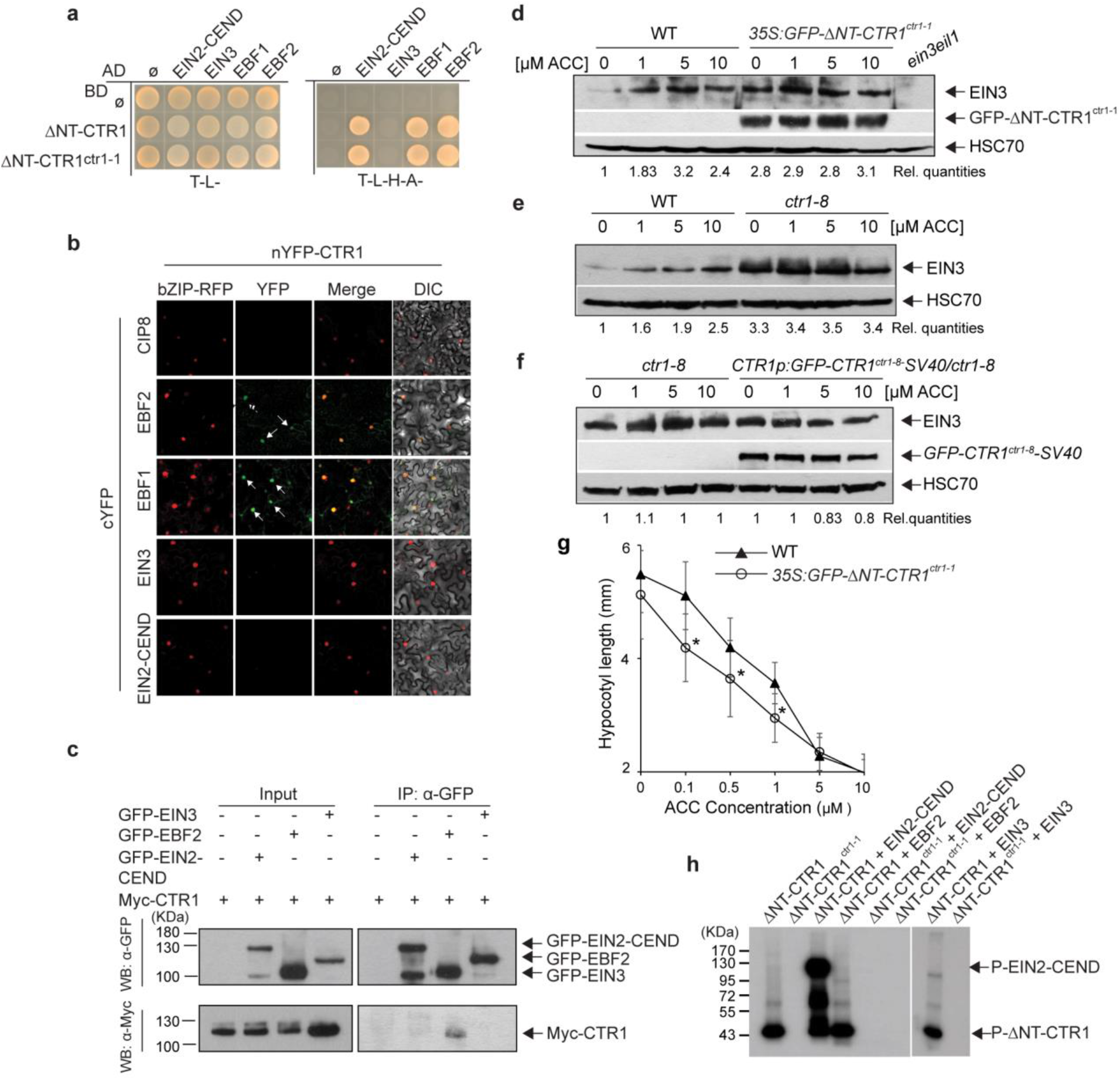
Nuclear-localized CTR1 stabilizes EIN3 via a non-catalytic function. **a**, The indicated bait and prey constructs were co-transformed into AH109, and the growth of the transformed AH109 was monitored on selection medium. **B,** BiFC assay for full-length WT CTR1 and nuclear ethylene signaling proteins in *N. benthamiana* in the presence of ACC. COP1-interacting protein 8 (CIP8) and bZIP-RFP were used as a negative control and a nuclear subcellular marker, respectively. **c**, Co-immunoprecipitation analysis of CTR1 and EBF2 in *N. benthamiana*. **d,** Seedlings expressing ΔNT-CTR1^ctr1-1^ had higher basal levels of EIN3 than WT seedlings. Three-day-old dark-grown seedlings were treated with the indicated concentrations of ACC for 2 h, and total protein extracts were used for immunoblotting with anti-EIN3 and HSC70 antibodies. **e, f,** *ctr1-8* (**e**) and seedlings expressing GFP-CTR1^ctr1-8^-SV40 NLS in the *ctr1-8* mutant (**f**) expressed higher basal levels of EIN3 than WT seedlings. Rel. quantities represent the ratio of the intensity of the EIN3 band to HSC70 band signals, and these values are expressed relative to the intensity of EIN3/HSC70 in the WT with no ACC treatment value, which was set to 1. **g,** Dark-grown *35:GFP-ΔNT-CTR1^ctr1-1^* seedlings have an enhanced ethylene response after exposure to the ethylene precursor ACC. ACC dose-response curves for the hypocotyl length of 3-d-old dark-grown WT and *35:GFP-ΔNT-CTR1^ctr1-1^* seedlings. Control treatments included no ACC. Error bars, SD (*n* ≥ 24). * *p* < 0.0001, Student’s *t*-test. **h**, CTR1 does not phosphorylate EBF2. *In vitro* kinase assay for purified ΔNT-CTR1 or ΔNT-CTR1^ctr1-1^ with EIN2-CEND, EIN3, or EBF2. EIN2-CEND and EIN3 were used as a positive and negative control, respectively. The indicated proteins were incubated together in kinase reaction buffer and then separated by SDS/PAGE, and the incorporated radiolabel was detected by autoradiography.

The ethylene response kinetics results showed that the enhanced nuclear localization of CTR1 delayed the restoration of seedling growth to basal levels after the removal of ethylene. Because EIN3 overexpression or a lack of EBF2 leads to similar recovery response kinetics, we examined endogenous EIN3 protein levels in the WT and seedlings overexpressing ΔNT-CTR1^ctr1-1^ (*35S:GFP-ΔNT-CTR1^ctr1-1^/Col-0*) in response to different ACC concentrations. In agreement with findings from prior studies^28, 29^, ACC stabilized EIN3, displaying an ACC dosedependent increase in the WT, whereas, *35S:GFP*-Δ*NT-CTR1^ctr1-1^/Col-0* seedlings expressed significantly higher basal levels of endogenous EIN3, which were not further stabilized by ACC (Fig. 5d). Likewise, *ctr1-8* and *CTR1p:GFP-CTR1^ctr1-8^-SV40/ctr1-8* seedlings, both of which showed delayed growth recovery after ethylene withdrawal (Fig. 4b), expressed higher basal levels of EIN3 than the WT seedlings, which did not show further EIN3 stabilization with higher ACC treatment (Fig. 5e and 5f). Consistent with the higher levels of endogenous EIN3 proteins, the hypocotyls of *35S:GFP-ΔNT-CTR1^ctr1-1^/Col-0* seedlings were significantly shorter than those of the WT over a broad range of ACC concentrations (Fig. 5g). *In vitro* kinase assays did not show any phosphorylation of EBF2 by CTR1 (Fig. 5h), corroborating the observation that CTR1 kinase activity is not involved in CTR1’s nuclear movement and the regulation of EIN3 protein stability. Together, these results demonstrate that, after translocation to the nucleus, CTR1 directly interacts with the EBFs and consequently inhibits the EBF-mediated degradation of EIN3.

## Discussion

After the perception of environmental signals, the specificity of cellular responses to stress is mediated by the spatial and temporal dynamics of downstream signaling networks. Here, we provide convincing evidence of a mechanism linking the spatiotemporal regulation of CTR1 to organismal ethylene responses. We discovered that, after inactivation by ethylene, CTR1 re-locates from the ER to the nucleus and subsequently enhances nuclear ethylene responses by stabilizing EIN3 by modulating EBF’s function without its kinase activity (Fig. 6).

**Fig. 6.**
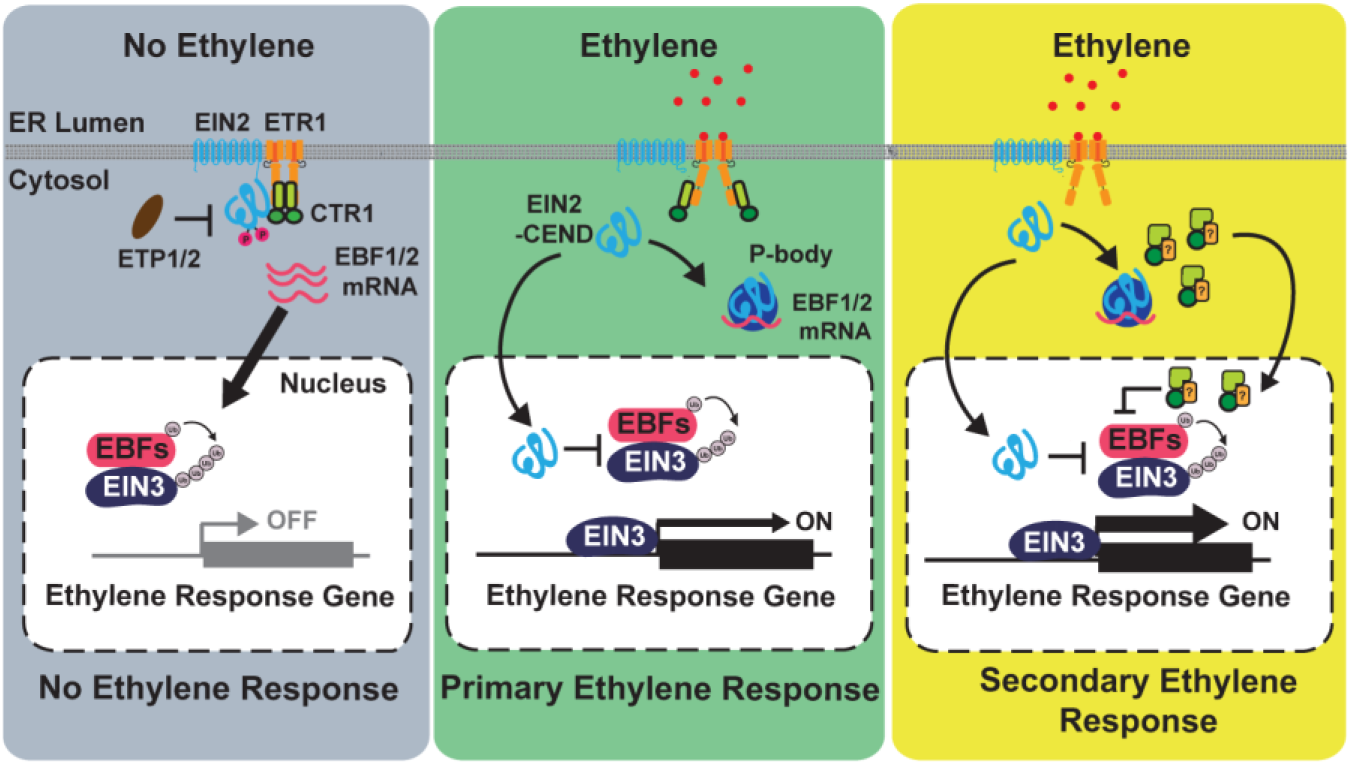
Model for ethylene-induced CTR1 nuclear translocation and stimulation of the ethylene response. In the absence of ethylene, the ethylene receptors, CTR1, and EIN2 mainly localize to the ER. Receptor-activated CTR1 phosphorylates EIN2, thus leading to the proteolytic degradation of EIN2 via EIN2-TARGETING PROTEIN 1 (ETP1) and ETP2. In the presence of ethylene, the receptors, and hence CTR1, are inactivated. Inactivated CTR1 no longer phosphorylates EIN2, and the C-terminal domain of EIN2 (EIN2-CEND) is proteolytically cleaved and translocates to the nucleus, where it initiates the primary ethylene response via EIN3 activation. EIN2-CEND is also targeted to processing (P-) bodies and suppresses EBF mRNA translation. After the nuclear translocation of EIN2-CEND, CTR1 is released from the receptors and relocates to the nucleus via an unknown mechanism. The nuclear-localized CTR1 directly interacts with EBF2, thus stabilizing EIN3 via an unknown mechanism and ultimately stimulating the ethylene response. The yellow square with a question mark indicates an unknown cargo protein that delivers CTR1 to the nucleus. Arrows indicate the movement of ethylene signaling components or positive influence; blunted ends indicate inhibition.

The failure of CTR1^ctr1-8^ protein to translocate to the nucleus was unpredicted, because weak interaction of CTR1^ctr1-8^ with ethylene receptors would increase the release of CTR1^ctr1-8^ from the ER, thus promoting nuclear translocation. Full-length WT CTR1 in the loss of function ethylene receptor mutants and CTR1 proteins with N-terminal deletion in the WT constitutively localized in the nucleus, suggesting that the interaction with ethylene receptors prevents CTR1 release from the ER. We speculated that the modification of CTR1 after ethylene binding to the receptors may relieve N-terminus-mediated inhibition of CTR1, thereby enabling CTR1 release from the ER and subsequent nuclear translocation. Given the weak interaction with the ethylene receptors, the CTR1^ctr1-8^ mutant protein may not undergo this conformational change and consequently fail to translocate to the nucleus. Alternatively, *ctr1-8* mutation might prevent the mutant CTR1^ctr1-8^ protein from interacting with an unknown cargo protein that transports CTR1 to the nucleus, given that CTR1 lacks canonical NLSs.

CTR1 forms a complex with EBFs and consequently influences EIN3 protein stability, although the detailed underlying mechanism remains elusive. Intriguingly, this process is independent of the catalytic function of CTR1. Many non-catalytic functions of protein kinases have been reported in yeast and mammalian systems^30, 33^, but not in plants, demonstrating that their extensive roles in biological processes as scaffolds, allosteric regulators, and molecular switches beyond phosphorylating proteins^30, 31, 34–36^. Given that CTR1 does not interact with EIN3, CTR1 might allosterically inactivate the activity of EBFs, thus inhibiting EIN3 degradation. Alternatively, CTR1 might decrease free nuclear EBF pools that interact with EIN3 through the formation of CTR1-EBF complexes.

Analysis of ethylene growth response kinetics has been instrumental in examining various ethylene mutants; however, the detailed mechanism underlying the recovery kinetics remains unknown. Previous studies have shown that increased abundance of EIN3 is the main factor underlying the delayed recovery kinetics after ethylene removal^19^. Our studies have revealed that nuclear-localized CTR1 plays a role in the growth recovery kinetics of *Arabidopsis* hypocotyls by increasing EIN3 stability via its interaction with EBFs. Given its negative roles in the ethylene response, the CTR1-mediated stimulation of the ethylene response in the nucleus was unforeseen; however, many proteins have dual functions in signal transduction pathways, and some perform opposite activities in the same biological process^37–40^. For example, in human memory control, AGAP3, an NMDA receptor-interacting signaling protein, not only enhances the activity of memory formation but also decreases the activity of memory formation, depending on the perception of different signals^37^. The CTR1-mediated stimulation of ethylene responses in the nucleus might also be an elegant way in which plants maximize ethylene responses by turning a negative regulator into a positive regulator, thereby enabling rapid acclimation to stresses.

CTR1 lacks canonical NLSs. How it translocates from the ER to the nucleus, and whether other ethylene-activated components might be involved in this process, remains to be determined. Furthermore, elucidation of the mechanism underlying CTR1-mediated EBF regulation and the identification of a nuclear pathway that promotes rapid growth recovery would provide more insights advancing the mechanistic understanding of ethylene response regulation. Ethylene interacts with a myriad of internal and external stimuli in regulating plant growth and stress responses. Thus, revisiting crosstalk with other pathways by considering the nuclear role of CTR1 should be of great interest.

## References and Notes

## Acknowledgments

We thank Alison Delong for critical reading of the manuscript, Eric Schaller for providing ethylene receptor mutants, and Purdue Imaging Facility for confocal microscope service. This work was supported by grants from NSF (MCB-1817286) and Ralph W. and Grace M Showalter Research Trust to GMY, and NSF (IOS-1856248) to JJK and B.M.B (MCB-1817304).

## Author contributions

GMY and JJK conceived the experiments; GMY planned and supervised experiments; HYL, DHS, HLP, CP conducted experiments except ethylene-response kinetics; AB and BMB conducted ethylene response kinetics analysis; GMY, JJK, BMB, HYL, DHS, HLP, and CP contributed to the interpretation of the data; GMY wrote the manuscript with support from JJK and BMB. HYL, DHS, and HLP equally contributed to the manuscript.

## Data and materials availability

All data are available in the main text or the supplementary materials.

## Materials and Methods

### Plant materials and growth conditions

*Arabidopsis thaliana* Col-0 was used as the WT reference throughout the study. All plants were grown in either long-day or short-day conditions at 22°C ± 2°C or in vitro on Murashige and Skoog (MS) basal medium supplemented with 0.8% plant agar (pH 5.7) in a continuous light chamber at 21°C. All plants used were homozygous or T2 lines. Homozygous transgenic lines were identified by the segregation of antibiotic resistance followed by the confirmation of protein expression via immunoblot analyses.

### CTR1 constructs and site-directed *in vitro* mutagenesis for *Arabidopsis* transformation

All molecular cloning was performed with the Gateway (Invitrogen) or infusion cloning (Takara Bio USA) strategies unless otherwise specified. Tables S1, S2, and S3 list all primers used for cloning and mutagenesis. To create the *CTR1p-GFP-gCTR1* construct, we PCR-amplified three separate overlapping fragments (CTR1 promoter, YFP, and the genomic fragment of CTR1). The 0.96 Kb CTR1 promoter and 4.7 kb full length CTR1 genomic fragment were amplified with Col-0 genomic DNA as a template, and the YFP coding sequences were amplified with the binary vector pEarleyGate 104. Three overlapping fragments were subsequently subjected to an infusion reaction with *Stu*I- and *Xba*I-digested pEarleyGate 104, thus yielding the GFP-full-length CTR1 clone in the pEarleyGate 104 backbone. Genomic *CTR1* fragment mutations (G354E *ctr1-8* or D694E *ctr1-1*) were introduced into the WT genomic fragment via PCR-based mutagenesis, and the resulting fragments were used for infusion reactions, as described above. *CTR1p-YFP-gCTR1* and its mutant variant constructs were also constructed as described above, except YFP was used instead of GFP. To construct *35S:GFP-CTR1, GFP-CTR1^ctr1-1^, GFP-ΔNT-CTR1*, *GFP-ΔNT-CTR1^ctr1-1^*, and *GFP-CTR1^ctr1-8^*, we cloned the coding sequences of full length CTR1 or its kinase domain into the pENTR entry vector and subsequently transferred them to the binary vector pSITE2CA. Mutations were introduced in the coding sequences of CTR1 in the pENTR vector, and the sequences were further transferred to pSITE2CA. To create the *CTR1p-GFP-CTR1^ctr1-8^-SV40 NLS* construct, we added *SV40 NLS* sequences to the *CTR1p-GFP-CTR1^ctr1-8^* fragment via overlapping PCR and cloned the resulting fragment into the pEarleyGate 104 vector via infusion as described above.

### Bimolecular fluorescence complementation constructs

The following coding sequences including a stop codon were transferred from the pENTR vector into pCL112 to generate N-terminal nYFP-fusions: *CTR1* and ENAP. The following coding sequences without a stop codon were transferred from the pENTR vector into *pBAT-YFPc* to generate C-terminal cYFP-fusions: *EIN3, EIN2-CEND*, and *CIP8*. The coding sequences of *EBF1* and *EBF2* in pENTR were transferred into pCL113, thus creating *cYFP-EBF1* and *cYFP-EBF2*.

### Phos-Tag gel analysis

Preparation of Phos-Tag polyacrylamide gels and subsequent immunoblotting were performed according to the manufacturer’s instructions. The Phos-Tag gel was prepared with 5 ml of 8% acrylamide and 20 μM Phos-Tag (Wako) for the resolving gel and 4 mL of 4.5% acrylamide for the stacking gel. After completion of electrophoresis, the gel was incubated in 100 mL transfer buffer containing 0.2% (w/v) SDS and 10 mM EDTA for 30 min. Protein was blotted onto nitrocellulose, which was then blocked with 5% nonfat milk. The membranes were then probed with a 1:5000 dilution of Roche anti-GFP (Sigma Aldrich Cat# 11814460001) or a 1:5000 dilution of anti-mouse HRP secondary antibody (ThermoFisher).

### Time-lapse growth recovery analysis

To measure the ethylene response growth kinetics of seedlings, we grew seedlings on vertically oriented Petri plates in the dark to a height of 3 to 4 mm (42–46 h) before the beginning of the growth-rate measurements. The agar plates were placed vertically in a holder and fitted with a lid for continuous gas flow (100 mL min^-1^). Seedlings were grown in air for 1 h, and ethylene (typically 1 or 10 ppm) was applied for 2 h and then removed. Images were acquired every 5 min with a CCD camera fitted with a close-focus lens with illumination provided by infrared LEDs. The growth rates of the hypocotyls in every time interval were then calculated. Under these conditions, the equilibration time of the chamber at these flow rates was approximately 30 sec, which was much faster than the image acquisition time. From the results, we determined how various mutations affected growth inhibition kinetics when ethylene was added and affected the recovery kinetics when it was removed.

### Yeast-two-hybrid assays

The coding sequences of full-length *CTR1* or its kinase domain (Δ*NT-CTR1*) with or without *ctr1-1* mutation (D694E) in the pENTR GW entry vector were transferred into pGBKT7 or pGADT7. The resulting bait clones were paired with *EIN2-CEND, EIN3, EBF1*, or *EBF2* in the pGBKT7 or pGADT7 vector, and their interactions were tested in yeast. Positive interactions between the prey and bait were selected on medium lacking histidine, tryptophan, leucine, and adenine. Because of the autoactivation of full-length *CTR1* in pGBKT7, we used *ΔNT-CTR1* or *ΔNT-CTR1^ctr1-1^* to generate *CTR1* bait constructs.

### Confocal microscopy

All imaging of GFP, YFP, and RFP was performed with a laser-scanning confocal microscope (Zeiss LSM880 upright). Samples were directly mounted on a glass slide in water. For imaging of *Arabidopsis* seedlings transformed with CTR1 constructs, the seedlings were grown on MS medium with or without 100 μM AgNO3 supplementation in the dark for 3 d. For ACC treatment, seedlings on MS without AgNO3 were treated with 200 μM ACC dissolved in water for 2 h. For ethylene treatment, seedlings were grown directly on MS medium in GC vials for 3 d. The GC vials were subsequently capped, injected with 1 ppm ethylene gas, and incubated for 2 h. For nuclear imaging, 3-d-old dark-grown seedlings were treated with 200 μM ACC for 2 h, incubated with Hoechst33342 solution (Invitrogen) for 30 min, and briefly washed before mounting on slides. For imaging light-grown seedlings, plants were grown on MS in constant light for 5 d. For imaging of fluorescence signals in protoplasts, transfected protoplasts were incubated with 200 μM ACC for 2 h in the dark and then examined. To image BiFC, leaf disks of infiltrated tobacco leaves with CTR1 and counterpart constructs (*EIN2-CEND, EIN3, EBF1, EBF2, ENAP1*, and *CIP8*) were mounted on glass slides in water.

### Coimmunoprecipitation assays

*N. benthamiana* leaves were infiltrated with agrobacteria co-transformed with the plasmids of interest and incubated for 3 d in a growth chamber. Total proteins were extracted from the infiltrated leaves and homogenized in co-immunoprecipitation buffer (25 mM Tris, pH 7.5, 150 mM NaCl, 5 mM EDTA, pH 8.0, 1 mM DTT, 1 mM PMSF, and 1× protease inhibitor cocktail). After quantification of the protein concentration with a Bradford assay, an equal amount of total protein extracts was incubated with anti-GFP (Sigma Aldrich) overnight at 4°C, then incubated with Protein A/G magnetic beads (Fisher) for 1 h at room temperature with gentle shaking. The total protein suspension containing the magnetic beads was applied to a magnetic column, washed three times with co-IP buffer, eluted with boiled 2× SDS sample buffer, and subjected to immunoblotting.

### Nuclear-cytoplasmic fractionation

Three-d-old etiolated Arabidopsis seedling was grinded with ice-cold lysis buffer (20 mM Tris, pH7.4, 25 % glycerol, 20 mM KCl, 2 mM EDTA, pH 8.0, 2.5 mM MgCl2, 250 mM sucrose, 40 μM MG132, 5 mM DTT, 1 mM PMSF, 1x Protease inhibitor cocktails) followed by filtration with 2 layers of Miracloth. The flow-through sample was collected and 200 μl was saved as total proteins. The flow-through was spun at 1,500g for 10 min to pellet the nuclei and the supernatant was transferred to new tubes and saved as the cytoplasmic fractions. The resulting pellet was resuspended in washing buffer (20 mM Tris, pH 7.4, 25% glycerol, 2.5 mM MgCl2, 25% Triton X-100, 20 μM MG132, 5 mM DTT, 1 mM PMSF, 1x Protease inhibitor cocktails) and spun at 1,500g for 10 min. After repeating washing the pellet with the washing buffer five times, the pellet was resuspended in the SDS sample buffer. The samples were resolved by SDS-PAGE and transferred to nitrocellulose membrane for immunoblotting analysis with anti-GFP (Sigma), −BIP (enzolifesciences), and −Histon H3 (Invitrogen) antibodies.

### In vitro kinase assays

A total of 20 ng purified His6-ΔNT-CTR1 or His6-ΔNT-CTR1^ctr1-1^ protein was incubated with 100 ng of His_6_-EIN2^WT^-His_6_, His_6_-EBF2, or His_6_-EIN3 in kinase reaction buffer [50 mM Tris (pH 7.5), 10 mM MgCl_2_, 1× Roche Complete Protease Inhibitor mixture, and 1 μCi [γ-^32^P] ATP] for 30 min at room temperature. After incubation, the reactions were terminated by boiling in 6× Laemmli SDS sample buffer for 3 min. Samples were subjected to SDS/PAGE, dried, and visualized by autoradiography.

### Real-time quantitative PCR analysis

Total RNA was prepared with RNeasy Plant Mini Kits (QIAGEN) and reverse transcribed with SuperScript II reverse transcriptase (Invitrogen) according to the manufacturers’ instructions. Quantitative RT-PCR was performed with PowerUP™SYBRGreen Master Mix (Applied Biosystems). The primers used are listed in Table S1. Three biological replicates were analyzed with three technical replicates per sample. The relative expression of candidate genes was normalized to *Actin 2*.

### Immunoblot analysis

Three-d-old dark-grown seedlings were treated with different concentrations of ACC for 2 h. The harvested seedlings were weighed, and 2× SDS sample buffer was added to the seedlings in proportion to their weight. The samples were then homogenized with a pestle. Subsequently, the same amount of total protein extract from each sample was boiled for 3 min and resolved through SDS-PAGE followed by immunoblotting with anti-EIN3 (Agrisera) or anti-HSC70 (Enzo Life Science). Signals were detected with SuperSignal West Pico or Femto Maximum Sensitivity Substrate (Thermo Fisher Scientific), and band intensities were measured in Image J software.

