## Supplementary information for "Ethylene-triggered subcellular trafficking of CTR1 enhances the response to ethylene gas"

**This PDF file includes:**

Supplementary Fig. 1-11
Supplementary Table 1

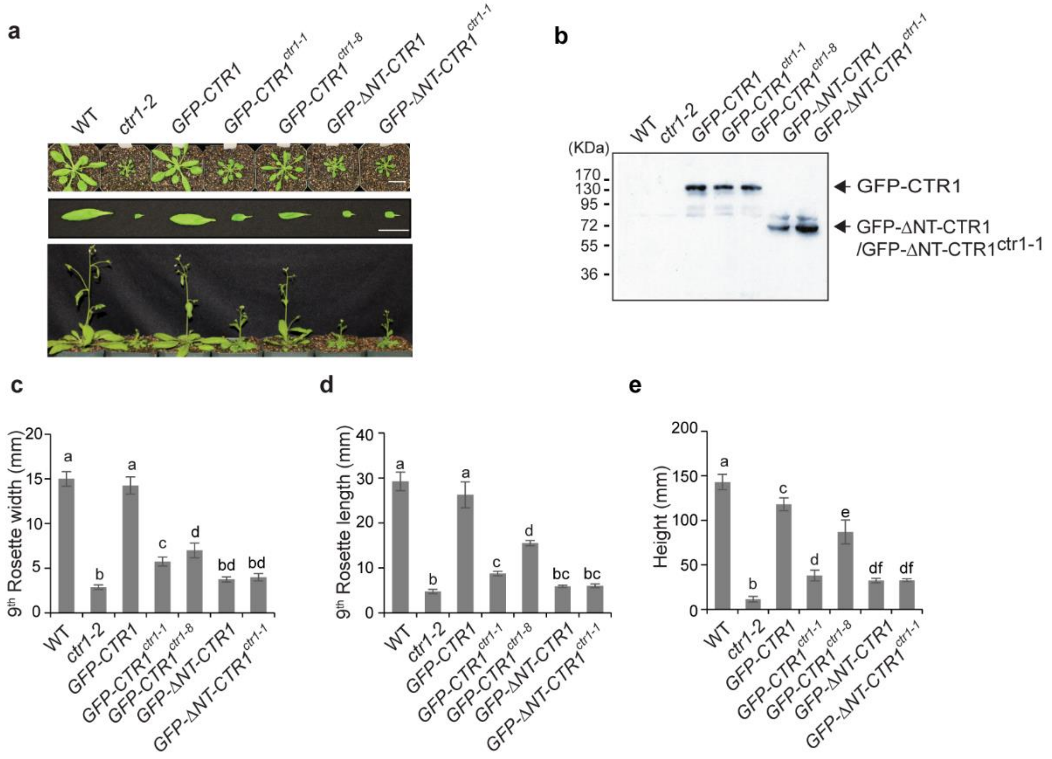


### Supplementary Figure 1

**Supplementary Figure 1. Rescue of *ctr1-2* by *CTR1* transgenes in light.**

**a**, Representative 2-week-old (top two rows) or 4-week-old (bottom row) light-grown WT and transgenic lines expressing various genomic *CTR1* transgenes from the native promoter in the *ctr1-2* background. Scale bar, 20 mm. **b**, Western blot analysis of GFP-fused CTR1 proteins expressed in seedlings in (**a**). **c-e**, Quantification of the width (**c**) and length(**d**) of the 9$^{th}$ rosette leaves of 2-week-old plants and the height (**e**) of 4-week-old plants in (**a**). Data represent the means and SD (*n*=4). Different letters indicate significant differences at $p < 0.05$ (one-way ANOVA, post-hoc Tukey's HSD). Data represent the means and SD (*n*=4).

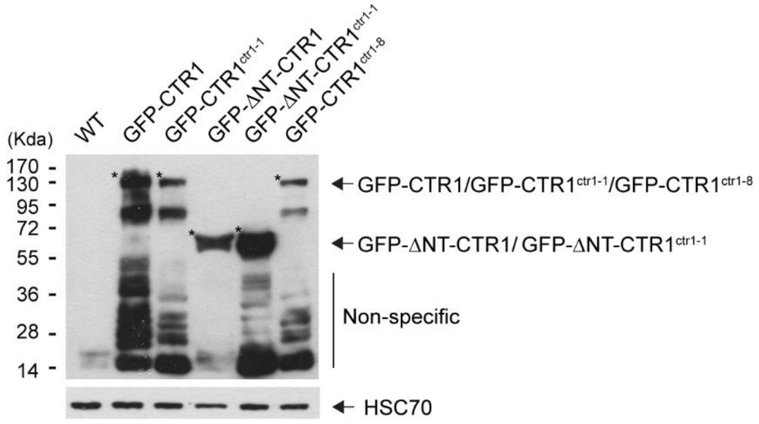


#### Supplementary Figure 2

**Supplementary Figure 2. Western blotting analysis of GFP-fused CTR1 in 35S promoter-driven overexpression lines.** The total protein extracts of 3-d-old dark-grown seedlings were subjected to immunoblotting analysis with anti-GFP and anti-HSC70. * indicates the corresponding protein bands.

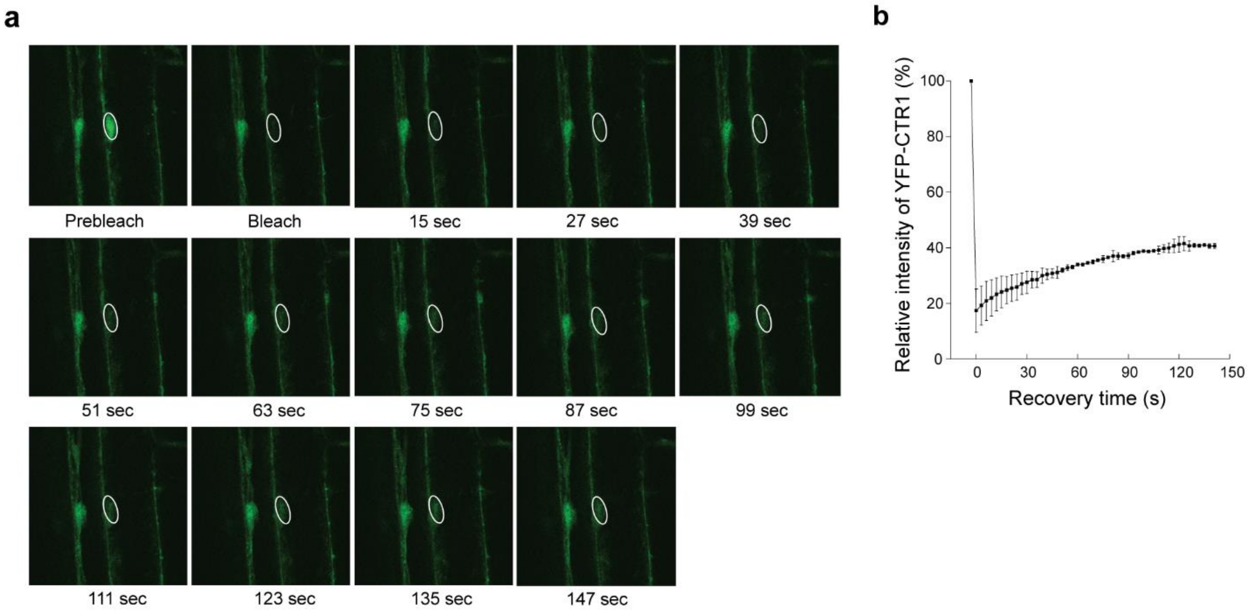


#### Supplementary Figure 3

**Supplementary Figure 3. Fluorescence recovery after photobleaching analysis of the GFP-CTR1 in the hypocotyls of *pCTR1:GFP-gCTR1/ctr1-2* seedlings. a,** Consecutive confocal images of GFP-CTR1 in hypocotyl nuclei before and after photobleaching. The photobleached area is indicated by white circles. **b**, Recovery kinetics of the circled nuclear area. After bleaching, images were taken for 3 seconds during recovery photobleaching. Error bar, SE.

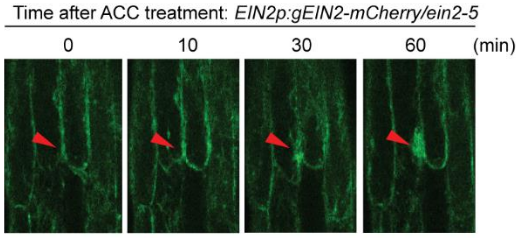


**Supplementary Figure 4**

**Supplementary Figure 4. Time-lapse analysis of EIN2 in the hypocotyls of dark-grown seedlings.** Time-lapse image series of hypocotyl cells expressing EIN2-mCherry in 3-d-old etiolated seedlings after exposure to 200 µM ACC, visualized by confocal microscopy. Arrows track specific cell nuclei, showing the accumulation of EIN2-mCherry in response to ACC.

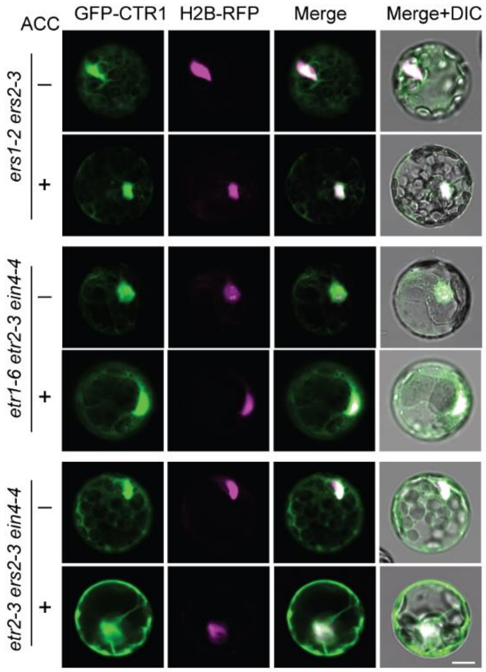


Supplementary Figure 5

**Supplementary Figure 5. Subcellular localization of CTR1 in various ethylene receptor mutants.** *Arabidopsis* mesophyll protoplasts from indicated high order ethylene receptor mutants were transfected with a *GFP-CTR1* plasmid, and the subcellular localization of GFP-CTR1 was observed by confocal microscopy. H2B-RFP is a nuclear marker protein. Scale bar, 10 µm.

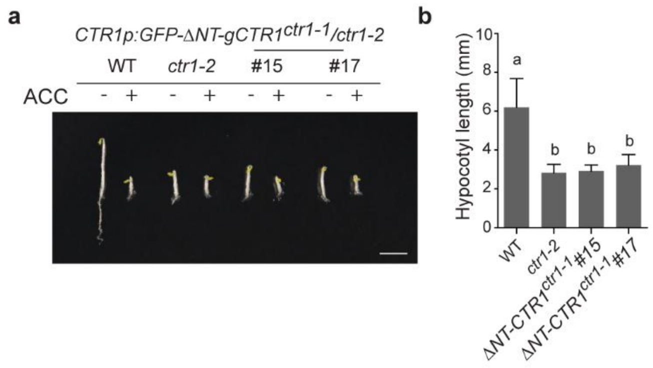


#### Supplementary Figure 6

**Supplementary Figure 6. The transgene encoding the CTR1 kinase domain without kinase activity does not rescue *ctr1-2*.**

**a**, Representative images of etiolated WT and seedlings expressing the CTR1 kinase domain only with *ctr1-1* mutation grown on MS medium with or without 10 μM ACC. Scale bar, 5 mm. **b**, Quantification of hypocotyl lengths of seedlings in (**a**) without ACC treatment. Different letters indicate significant differences at $p < 0.001$ (one-way ANOVA, post-hoc Tukey's HSD). Data represent the means and SD; $n \geq 35$.

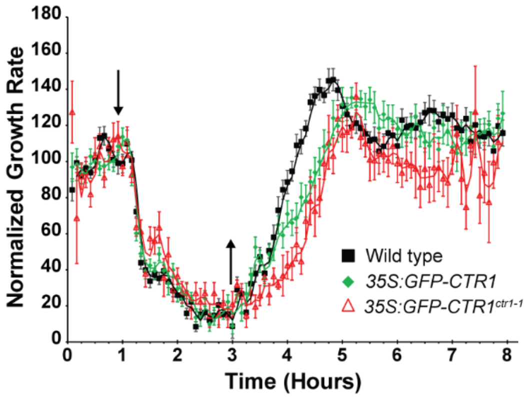


#### Supplementary Figure 7

**Supplementary Figure 7. Nuclear-localized CTR1 modulates the growth recovery of hypocotyls after ethylene removal.** The hypocotyl growth rate in response to ethylene was recorded for 1 h in air, followed by 2 h exposure to 10 ppm ethylene, and then a 5 h recovery in air. Ethylene was introduced 1 h after measurements were initiated (down arrow) and then removed 2 h later (up arrow). The ethylene growth responses of WT, *35S:GFP-CTR1*, and *35:GFP-CTR1*$^{ctr1\text{-}1}$ are shown in the graph. Data were normalized to the growth rate in air before treatment with ethylene. Error bars indicate SE (*n* ≥ 6). The experiments were repeated at least three times and generated similar results.

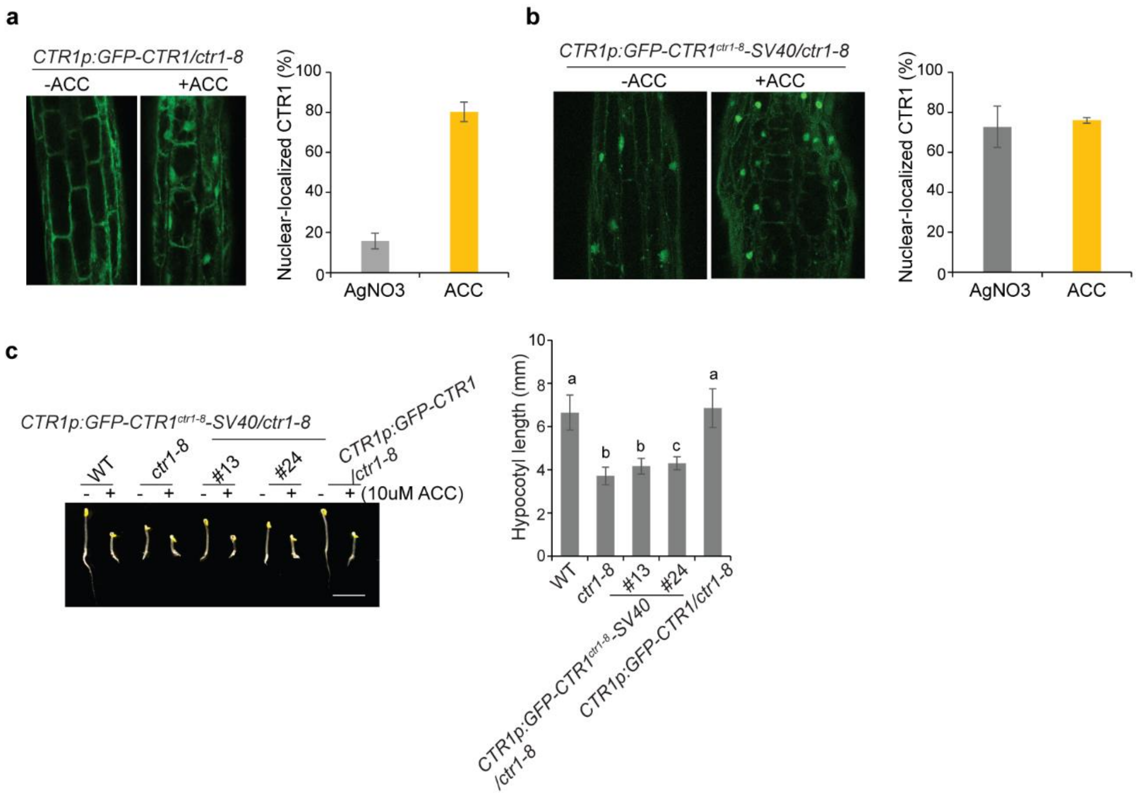


### Supplemental Figure 8

**Supplementary Figure 8. Constitutive nuclear localization of GFP-CTR1$^{ctr1\text{-}8}$-SV40 NLS.**
**a-b**, Three-d-old dark-grown seedlings were treated with or without 200 μM ACC for 2 h and the localization of GFP-CTR1 was imaged. The graphs represent the ratio of nuclear-localized CTR1 in dark-grown seedlings. Error bars indicate SD ($n \geq 3$). **c**, The WT *GFP-CTR1* transgene, but not *GFP-CTR1*$^{ctr1\text{-}8}$*-SV40*, fully rescued *ctr1-8*. Seedlings were grown for 3 d in the dark with or without ACC or ethylene. The graph represents the quantification of hypocotyl lengths of seedlings grown on growth medium without ACC. Different letters indicate significant differences at $p < 0.05$ (one-way ANOVA, post-hoc Tukey's HSD). Data represent the means and SD ($n >$ 22). Scale bar, 5 mm.

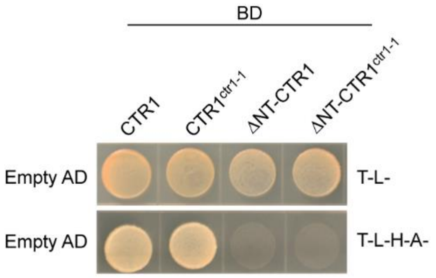


#### Supplementary Figure 9

**Supplementary Figure 9. Full-length CTR1-fused to a DNA-binding domain is autoactivated in the yeast-2-hybrid assay.** The coding regions of the full-length CTR1 or N-terminally deleted CTR1 with or without kinase activity were fused to a DNA-binding domain (bait). AH109 yeast strains expressing bait and empty prey plasmids were grown on selection medium.

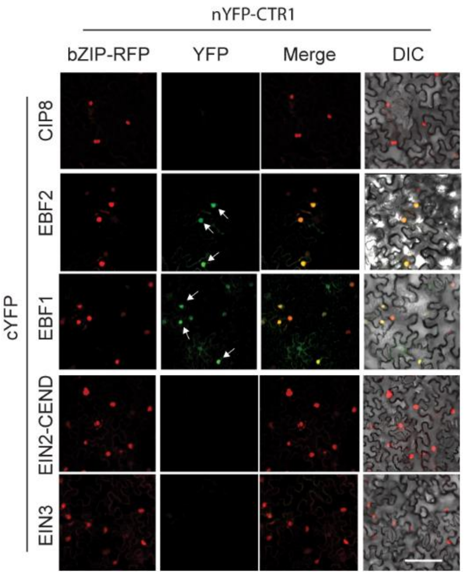


**Supplementary Figure 10**

**Supplementary Figure 10. CTR1 interacts with EBF1 or EBF2 in the nucleus in the absence of ACC.** BiFC assay for full-length WT CTR1 and nuclear ethylene signaling proteins in *N. benthamiana* in the absence of ACC. Tobacco leaves were infiltrated with agrobacteria transformed with the indicated constructs along with the bZIP-RFP nuclear marker, and further incubated for 3 d before imaging of protein-protein interactions with confocal microscopy. CIP8, COP1-interacting protein 8, was used as a negative control. Scale bar, 100 μm.

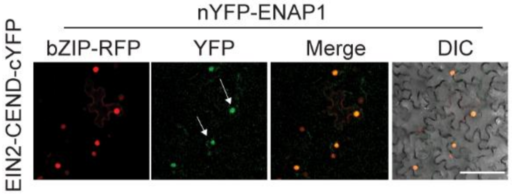


#### Supplementary Figure 11

**Supplementary Figure 11. EIN2-CEND interacts with ENAP1 in the nucleus**.
*N. benthamiana* was infiltrated with agrobacteria co-transformed with EIN2-CEND-cYFP, nYFP-ENAP1, and the bZIP-RFP nuclear marker, and incubated for 3 d before imaging. Arrows indicate the reconstituted YFP signals. Scale bar, 100 μm.
